# Coordinate based meta-analysis of networks in neuroimaging studies

**DOI:** 10.1101/407270

**Authors:** CR Tench, Radu Tanasescu, CS Constantinescu, DP Auer, WJ Cottam

## Abstract

Meta-analysis of published neuroimaging results is commonly performed using coordinate based meta-analysis (CBMA). Most commonly CBMA algorithms detect spatial clustering of reported coordinates across multiple studies by assuming that results relating to the common hypothesis fall in similar anatomical locations. The null hypothesis is that studies report uncorrelated results, which is simulated by random coordinates. It is assumed that multiple clusters are independent yet it is likely that multiple results reported per study are not, and in fact represent a network effect. Here the multiple reported effect sizes (reported peak Z scores) are assumed multivariate normal, and maximum likelihood used to estimate the parameters of the covariance matrix. The hypothesis is that the effect sizes are correlated. The parameters are covariance of effect size, considered as edges of a network, while clusters are considered as nodes. In this way coordinate based meta-analysis of networks (CBMAN) estimates a network of reported meta-effects, rather than multiple independent effects (clusters).

CBMAN uses only the same data as CBMA, yet produces extra information in terms of the correlation between clusters. Here it is validated on numerically simulated data, and demonstrated on real data used previously to demonstrate CBMA. The CBMA and CBMAN clusters are similar, despite the very different hypothesis.

## Introduction

Coordinate based meta-analysis (CBMA) is an approach commonly used to estimate the reliably observable effects from multiple independent, but related by a common hypothesis, neuroimaging studies [1-10]. It is employed to meta-analyse (amongst others) voxel based morphometry (VBM), functional magnetic resonance imaging (fMRI), and functional positron emission tomography (PET) studies, and uses only the reported summary statistics; coordinates and/or Z scores. The results should be interpreted as the distribution of activation peaks (foci) rather than estimates of whole voxel-wise activation (in the case of fMRI) clusters [1, 4, 10].

The most popular method of performing CBMA is the activation likelihood estimate (ALE) algorithm [1, 3, 5, 6, 9, 11]; other kernel based methods have also been described, but having quite similar approaches. The approach is to smooth, using a Gaussian kernel of ~10mm (conditional on the number of subjects) full width half max (FWHM), the reported foci to form a modelled activation image for each independent study, which are subsequently combined into a single ALE map. A permutation test is performed to establish an ALE threshold that controls the type 1 error rate in a principled manner; this test involves replacing the coordinates by random coordinates representing studies that are uncorrelated. Post threshold, isolated voxel clusters are formed representing an estimate of the distribution of reported foci relating to the common hypothesis. It is these clusters and their anatomical locations that form the output of CBMA algorithms and the result on which the interpretation and conclusion are based. Recently a coordinate based random effect size (CBRES) meta-analysis method has been reported [10]. CBRES differs from other kernel based methods in that cluster forming happens prior to significance testing using a method based on the popular DBSCAN (Density-based spatial clustering of applications with noise) algorithm [12], and these clusters are of foci rather than voxels. The algorithm then performs a conventional meta-analysis of reported effect sizes in each cluster; the reported effect sizes are the reported peak Z scores standardised by the study sample size. The use of effect sizes has the benefit of bringing standard meta-analysis methodology to CBMA, and also making coordinate based meta-regression possible.

CBRES estimates a meta-effect size in each cluster by fitting, using maximum likelihood, a univariate normal distribution parameterised with a mean effect and a random effect variance to account for differences between studies. Generally multiple effects are reported per study, each with an associated effect size (Z score or t statistic), and therefore univariate normal distributions need to be fitted to multiple clusters. Use of the univariate normal implies an assumption of independence of the clusters, yet it is likely they represent a spatial pattern of correlated results; a strong activation (large Z score) in one brain region might be related to a strong activation in another region. This assumption reflects the concept of the brain operating on a network level. In general CBMA does not consider dependence between spatially separated effects, and any correlation, or lack thereof, does not influence the results. However, using CBRES it has been demonstrated that effect sizes in multiple clusters can be correlated [13], and if these clusters are considered nodes of a network, and the correlation between the clusters as edges, then coordinate based metaanalysis of networks (CBMAN) becomes possible.

Coordinate based meta-analysis of networks uses the same data, coordinates and peak Z scores, as CBRES MA so both analyses can be performed simultaneously. Analysis proceeds by forming clusters from the reported coordinates and associated effect sizes just as in CBRES meta-analysis. Then, rather than univariate normal distributions fitted to effect sizes in each cluster, a multivariate normal (MVN) distribution is fitted to effect sizes in all clusters. The MVN distribution is parameterised such that there are means and variances (just as for CBRES MA) for each cluster, plus correlations relating the effect sizes between the clusters. It is these correlations that form the edges in the network of nodes (clusters).

This article describes coordinate based meta-analysis of networks. The methods involved in fitting MVN distributions are detailed, including the subtleties involved with censored data; summary results from neuroimaging studies are censored since only peak results exceeding a minimum threshold are reported. Ability to estimate correlation between censored effects is demonstrated by numerical experiments. Simulated networks are used to demonstrate the algorithm and real data from thermal painful stimulus fMRI studies, and VBM studies of multiple sclerosis, employed to show feasibility. Type 1 error rate control is by false discovery rate (FDR) [14]. The software to perform CBMAN is provided to use freely (search NeuRoi), and test experiments provided for validation.

## Methods

### The cluster forming algorithm

The method by which clusters are formed in CBMAN is based on the concepts of the popular DBSCAN algorithm [12]. DBSCAN aims to separate clusters of high density from surrounding low density noise. DBSCAN requires two parameters: a maximum clustering distance (MaxCD), and a minimum number of studies (MinStudies) contributing coordinates to the cluster. The MinStudies suggested for DBSCAN is at least the dimensionality plus one, and four is used for the three dimensional clustering problem considered here; coordinates from three studies forms only a surface, rather than a volume, in three dimensions. The purpose of the MaxCD is to allow formation of clusters where the spatial density of reported experimental results is high, while preventing cluster formation where experiments report spatially sparse results. The definition of the maximum clustering distance has here been updated from the definition presented in [10] to improve interpretability and ease of computation and does not substantially alter the clustering distance. In order to determine the MaxCD the coordinates are randomised as described in [10]. Then the proportion of coordinates involved in clusters (coordinates from at least four independent experiments fall within MaxCD of each other) computed. Sufficient iterations of this process are performed to accurately estimate this proportion. The MaxCD is then defined as that which results in only 5%, on average, of random coordinates forming clusters; 95% not clustered. This constraint imposes the requirement that the noise coordinates, as simulated by random coordinates, are unlikely to form clusters on average. An important feature of this approach is that it adapts the clustering distance to the number of studies included in the analysis; the larger the number of studies, the smaller the necessary clustering distance due to the increased density of reported coordinates. This is one of the major advantages of CBRES over other kernel based CBMA methods where a fixed size kernal results in a type 1 error rate that paradoxically increases with number of studies, and cluster growth that relates to the number of studies rather than being determined by the hypotheses tested by the studies.

For the clustering algorithm considered here, several deviations from DBSCAN have been implemented, each of which aim to improve resolution of clusters that overlap spatially. The first constrains the number of coordinates from independent studies that are overlapping (falling within the clustering distance of each other) to be reducing (but not strictly) away from the coordinate with the maximum overlap (the peak of the cluster).

The second modification is motivated by the OPTICS (ordering points to identify the clustering structure) algorithm [15], which forms part of a generalised DBSCAN algorithm. The modification allows for variable clustering distance (CD ≤ MaxCD) so that where coordinates reported by independent experiments are spatially very dense the CD is reduced, limiting the possibility of merging overlapping clusters. The choice of CD for a coordinate is the distance that envelopes coordinates from three other experiments; making the required MinStudies=4 overlapping experiments in total. This CD is then subject to a minimum value to prevent clusters fragmenting due to within cluster peaks that result only from the noise of sampling. This minimum can be estimated given the characteristics of a typical cluster, as detailed in [9]. To proceed, a number of coordinates equal to the number of independent studies is randomly generated from a trivariate normal distribution with diagonal covariance matrix with equal standard deviations 3.5mm; the distribution of coordinates then approximates that observed empirically. Because of the random nature it is possible there are multiple peaks within this cluster, which could cause the cluster to be fragmented by the clustering algorithm. By sampling the incidence of fragmentation (counting the proportion of times more than 1 cluster is detected by the clustering algorithm), a minimum clustering distance is deduced such that fragmenting occurs in very few (1%) of such clusters.

The final modification to DBSCAN is the requirement that coordinates are assigned to the nearest peak. It is possible that coordinates can potentially belong to multiple clusters, recruited by multiple peaks. The decision of cluster assignment is then based on the nearest peak. This is implemented as a shortest path distance in the clustering algorithm.

The pseudo code for the clustering algorithm is provided in the appendix.

### Standardised effect size

The effect sizes reported by most studies are the peak *Z* score; if *t* statistics or uncorrected p-values are reported instead, these can be converted to *Z* scores. This score depends on the number of subjects in the study, so may deviate considerably from being normally distributed, which is undesirable when fitting the MVN distributions used in CBMAN. The effect size (ES) used in CBMAN is the same as that used in CBRES [10], which is standardised by the number of subjects using

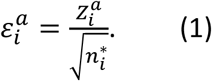

This specifies the effect size ɛ from study *i* in cluster *a* as a *Z* score standardised using the number of subjects *n*^*\**^ in study *i*. Just as in CBRES the number of subjects depends in whether the study is of a single group or a comparison between two groups. For single study groups the value of *n*^*\**^ is the number of subjects. For two group comparison studies the standardiser is

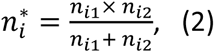

where *n*_*i1*_ is the number of subjects in group 1 and *n*_*i2*_ is the number of subjects in group 2.

### Fitting multivariate normal distributions to standardised effect sizes

The general form of the *k* dimensional multivariate normal distribution is

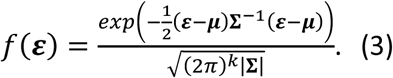

For *k* clusters discovered by the clustering algorithm the network has *k* nodes and the assumption of CBMAN is that the standardised reported effect sizes, ɛ, are distributed as specified in equation (3); this highlights a fundamental difference to CBRES meta-analysis, where effect sizes are considered univariate normal in *k* independent clusters. The network parameters, to be estimated by maximum likelihood, are then: μa column vector of means with *k* dimensions, and *∑* a symmetric covariance matrix of size *k*×*k*. Estimating the parameters of the MVN distribution can be a high dimensional problem if *k* is large; and as will be shown, a nonlinear one. Fortunately it is a property of the MVN that marginal distributions over subsets of the dimensions are themselves MVN distributions with the same subset of parameters. In CBMAN this fact is utilised to estimate parameters by fitting bivariate normal (BVN) distributions to pairwise clusters, reducing the problem to estimating multiple sets of just 5 parameters.

To fit BVN distributions to the standardised effect sizes in pairwise clusters at least two clusters must form. The BVN has five parameters to estimate by using maximum likelihood estimation (MLE): two means (*μ*) and two standard deviations (σ) plus a correlation (ρ). Effect size pairs for study *i* in clusters *a* & *b* are distributed as

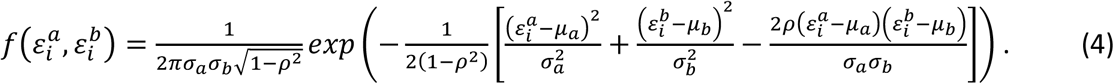

To estimate the parameters the log likelihood (LL) is maximised. There is one additive term to LL for each study in the analysis. A subtlety in the calculation is that some studies may not contribute a coordinate, and associated ES, to either or both of the clusters. In this instance the effect size is considered to be interval censored; it is known only that the value does not exceed a threshold level, and it is assumed that the censored value is drawn from the BVN distribution. Statistically the contribution to the LL of interval censoring is computed by integrating over regions/lines of the BVN distribution; this is why the problem of fitting the MVN distribution in CBMAN is non-linear.

When the ES from a study is not censored (the study contributes a coordinate to both clusters) the additive contribution to the LL is just the log of equation (4).

If the study contributes to only one of the clusters, say cluster *b*, one effect size is censored and known only to fall between ±α, where α is derived from the study *Z* score threshold for significance using equation (1); the threshold *Z* score is often reported, but is estimated by the minimum reported *Z* score otherwise. Then the contribution to LL is computed by integration of equation (4),

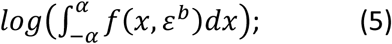

and similar if censored in cluster *b*. The integral in equation (5) can be computed analytically using the error function and the conditional distribution for the ES in cluster *a* given the ES in cluster *b*, however for this report it is computed numerically [16] using Simpson’s rule.

Another scenario considered by CBMAN is when a study contributes a coordinate to neither cluster. In this case both effect sizes are interval censored and known only to fall between thresholds ± α. The contribution to LL is then

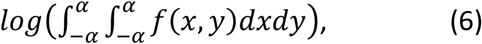

which is also computed numerically.

There is one other type of censoring common in neuroimaging studies whereby coordinates are listed but no ES is reported; such studies report only if the ES is positive or negative. The effect sizes are then left or right censored, known only to be less than *-α* (left censored) or greater than α (right censored). This type of censoring can be considered by integration over regions of the BVN distribution just as for interval censoring. The general LL term is

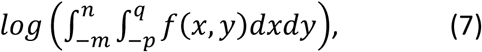

where the integral limits depend on whether censoring is left or right, or interval if the study reports no coordinate within the cluster. Table 1 indicates the limits for each scenario. Integrals with these limits are computed numerically, with ∞ limits approximated by α +6; this is around 5 or 6 standard deviations beyond the typical effect size and increasing this has been found not to alter the results in any significant way.

**Table 1.** Integral limits for the calculation, using equation (7), of the LL contribution for studies that do not report *Z* scores, *t* statistics, or uncorrected p-values.

| Censor type | Lower integral limit | Upper integral limit |
| --- | --- | --- |
| Left | $-\infty$ | $-\alpha$ |
| Right | $\alpha$ | ∞ |
| Interval | $-\alpha$ | $+\alpha$ |

### Thresholding the correlation between clusters

The correlation of effect sizes (edges) between clusters (nodes), estimated by maximum likelihood, determines the network in CBMAN. This produces a fully connected network (every node connected to every other node), which is not very revealing. A method of thresholding the edges is required to reduce the network such that the connections between nodes are meaningful by some criteria. In network analysis of the brain this thresholding is still an open question, but requiring correlation to be above a minimum magnitude is a common strategy [17]. In CBMAN there are often relatively few studies (10’s) so while the point estimate of the correlation could be high, the variance on the estimate could also be large; in this case thresholding by correlation alone may not be sufficient. The first threshold applied by CBMAN relates to the fact that correlation computed with just two points is not very meaningful, and with only three points there are few degrees of freedom to estimate the variance of the correlation. Without being too restrictive, CBMAN considers correlation between two clusters only if at least four studies contribute uncensored effect sizes to both clusters; censored effect sizes then either support or refute the correlation. Furthermore, as well as the option to set a minimum correlation CBMAN performs statistical hypothesis testing to identify edges for which there is statistical evidence.

For each edge there is a correlation of ES between the two clusters connected by the edge. This correlation is estimated by the maximum likelihood fit of the BVN model specified in equation (4). Evidence for a statistically significant correlation is obtained using the likelihood ratio test. This requires a null hypothesis where, in this case, the correlation parameter ρ is zero; for this hypothesis the parameters in equation (4) are fitted by maximum likelihood, but with the correlation parameter set to zero. The likelihood ratio test is performed by computing the LL for the alternative hypothesis (LL_A_; *ρ*not assumed to be zero) and also for the null hypothesis (LL_0_; ρ=0), then computing the random variable

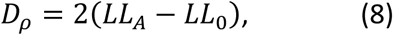

which is asymptotically chi-squared (χ ^2^) distributed with one degree of freedom. The p-value is then 1-q, where q is the quantile for *D*_*ρ*_ in the (χ^2^distribution.

The number of edges is quadratic in the number of nodes in the network. Therefore, the necessary correction for multiple statistical tests can escalate quickly. The purpose of the correlation threshold is to remove correlations that are too small to be important, and thereby reduce the number of tests and the multiple testing problem. In CBMAN a default threshold is computed by estimating the correlation coefficient that could be detected with 80% power, p-value<0.05, using Pearson’s correlation given the number of studies in the analysis. Statistical thresholding is then by FDR on p-values obtained by the likelihood ratio test only where the correlation exceeds the threshold.

### Other models

The hypothesis that the effect sizes are correlated between clusters is only one of the possibilities that can be tested. Just like CBMA it is also possible to compare two groups by modification of the parameters estimated by MLE. For example each group could have a different correlation, or different mean and correlation. Statistical comparison of coordinate based networks between the groups is then possible again using the likelihood ratio test, highlighting any difference in ES correlation. These models are not considered in this report, but are currently being developed and tested.

### Experiments

In this report the concept and computation are validated using simulated data. To demonstrate applied utility, coordinates extracted from published studies are used.

#### Simulated experiments

Numerical simulation is used to validate the code and to test: the validity of the asymptotic assumption of the likelihood ratio test, the ability to estimate parameters of the BVN distribution when data are censored, and the ability to identify known networks.

To test parameter estimation for censored data 50 BVN data are simulated with random parameters *μ*_*a*_, *μ*_*b*_, *σ*_a_, *σ*_b_*,ρ*; mean parameters are uniform random 0.6≤ *μ*≤1.2, standard deviations are uniform random 0.2≤*σ* ≤0.6, and correlation is uniform between ±1; these settings represent the range of results presented in [13]. Data are interval censored if they are <3.09/√20=0.7 (representing the ES minimum for commonly used Z score threshold of 3.09 and 20 subjects), and 10% of studies are left/right censored. The known simulation parameters and the corresponding estimates are qualitatively compared by scatter plot for examples with 20 studies and 50 studies.

To test the assumption that equation (8) is χ^2^ distributed with 1 degree of freedom under the null hypothesis 1000 simulated BVN data (using equation (4)) are generated with parameters *μ*_*a*_=*μ*_*b*_=1.0, σ_a_ =σ_b_ =0.5, and *ρ*=0; these settings represent typical results presented in [13]. The parameters are estimated by maximum likelihood under both alternative and null hypotheses and *D*_*ρ*_ computed using equation (8). The distribution of *D*_*ρ*_ is then qualitatively compared to the χ^2^ distribution for 10 studies and 20 studies and assuming 20 subjects and the commonly used Z score threshold of 3.09.

A final simulated experiment involves 6 clusters and four different covariance matrices generating four different networks. For these experiments the effect sizes are simulated with a mean Z score of 5, and a standard deviation of 1. Censoring is at a Z score of 3.09, the number of subjects is 20, and the number of studies 30. The correlation parameter is set at a moderate level of 0.6, and no threshold is applied before correcting for multiple statistical tests using FDR=0.05. The first network simulated involves a diagonal covariance matrix, so no network is expected since the correlations should be statistically zero; this is an important example as no network should be produced providing the correction for multiple tests is successful. A second network is simulated with covariance matrix having only one off diagonal covariance for which the expected network has two nodes and one edge. A third network is simulated with a block diagonal covariance matrix generating two independent networks. Finally a covariance matrix with all off diagonal elements involving the correlation is expected to produce a fully connected network.

#### Real data example: fMRI of thermally induced painful stimulation

An example of a functional MRI meta-analysis, painful thermal stimulation, was provided in the CBRES MA report. Here that data are used for CBMAN.

#### Real data example: Voxel based morphometry of Multiple Sclerosis

It is well known that multiple sclerosis causes atrophy of the grey matter, and repeatable patterns of atrophy have been demonstrated by multiple VBM studies. The data used in this example are taken directly from the CBRES report. Both CBRES meta-analysis and CBMAN are performed.

## Results

### Simulated data experiment

There are two requirements that need to be met for statistically valid meta-analysis of networks. Firstly, accurate estimates of correlation of ES between clusters must be possible for a reasonable number of studies; typically CBMA involves 10’s of studies. For simulated BVN distributed data, using realistic effect sizes and censoring thresholds, figure 1 shows error due to maximum likelihood estimation of parameters. With 20 studies (top) the parameters can be estimated, but as expected the sampling error is evident. Nevertheless, simulations of strong correlation are quite reliable. Increasing to 50 studies considerably improves estimates and therefore reliability of the analysis.

**Figure 1.**
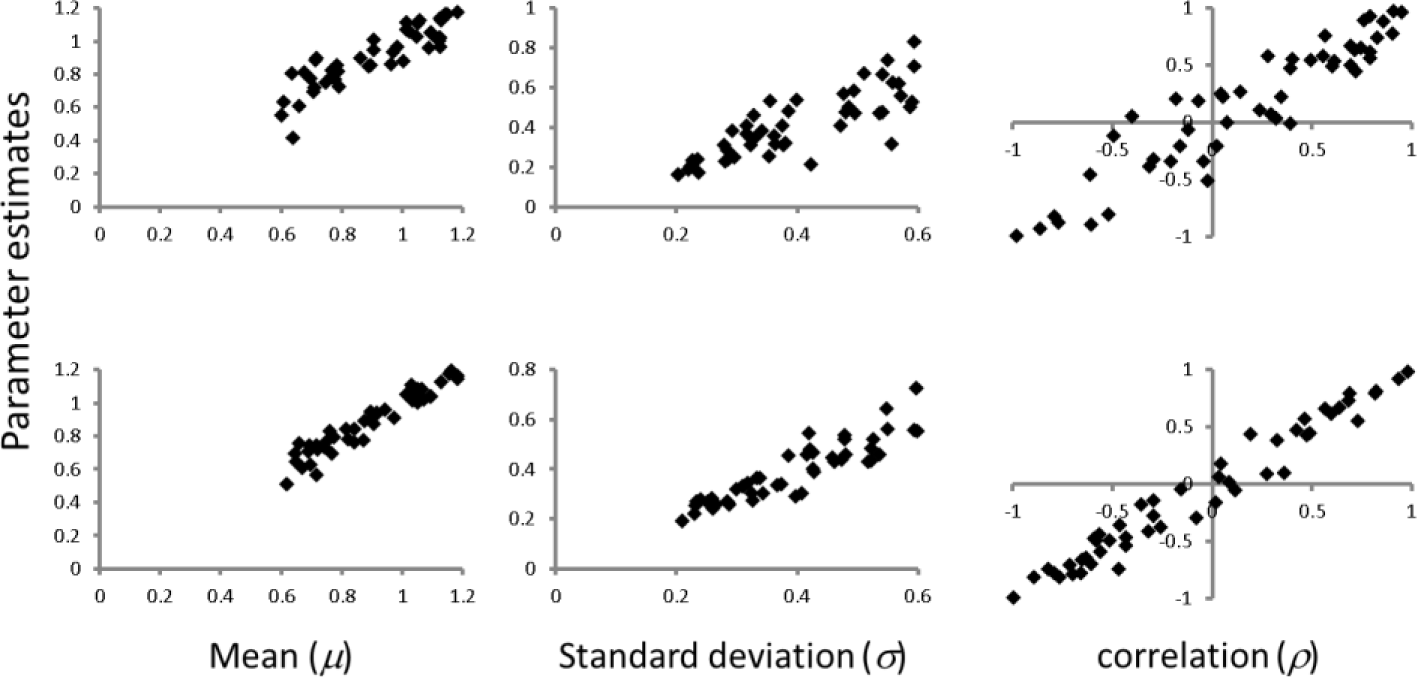
Demonstration of parameter estimation using simulated censored data. The top row is an example with 20 simulated studies, representing a small meta-analysis. The bottom row uses 50 simulated studies, representing a moderate sized meta-analysis. Estimates are better for the larger number of studies, as expected.

The second requirement is accurate assessment of statistical significance. CBMAN uses the likelihood ratio test to deduce p-values, but this is only valid asymptotically. It is therefore important that the null distribution of the test statistic (equation (8)) converge to its theoretical form (χ^2^ with 1 degree of freedom) with an appropriate number of studies. Figure 2 demonstrates that with only 10 studies this requirement is not met, and analysis might be liberal due to an excess of unexpectedly high test statistic under the null hypothesis; 10 studies represents a very small MA, but if performed a conservative FDR correction might be best. By 20 studies the test statistic has converged towards the χ^2^ distribution, validating the likelihood ratio test for CBMAN.

**Figure 2.**
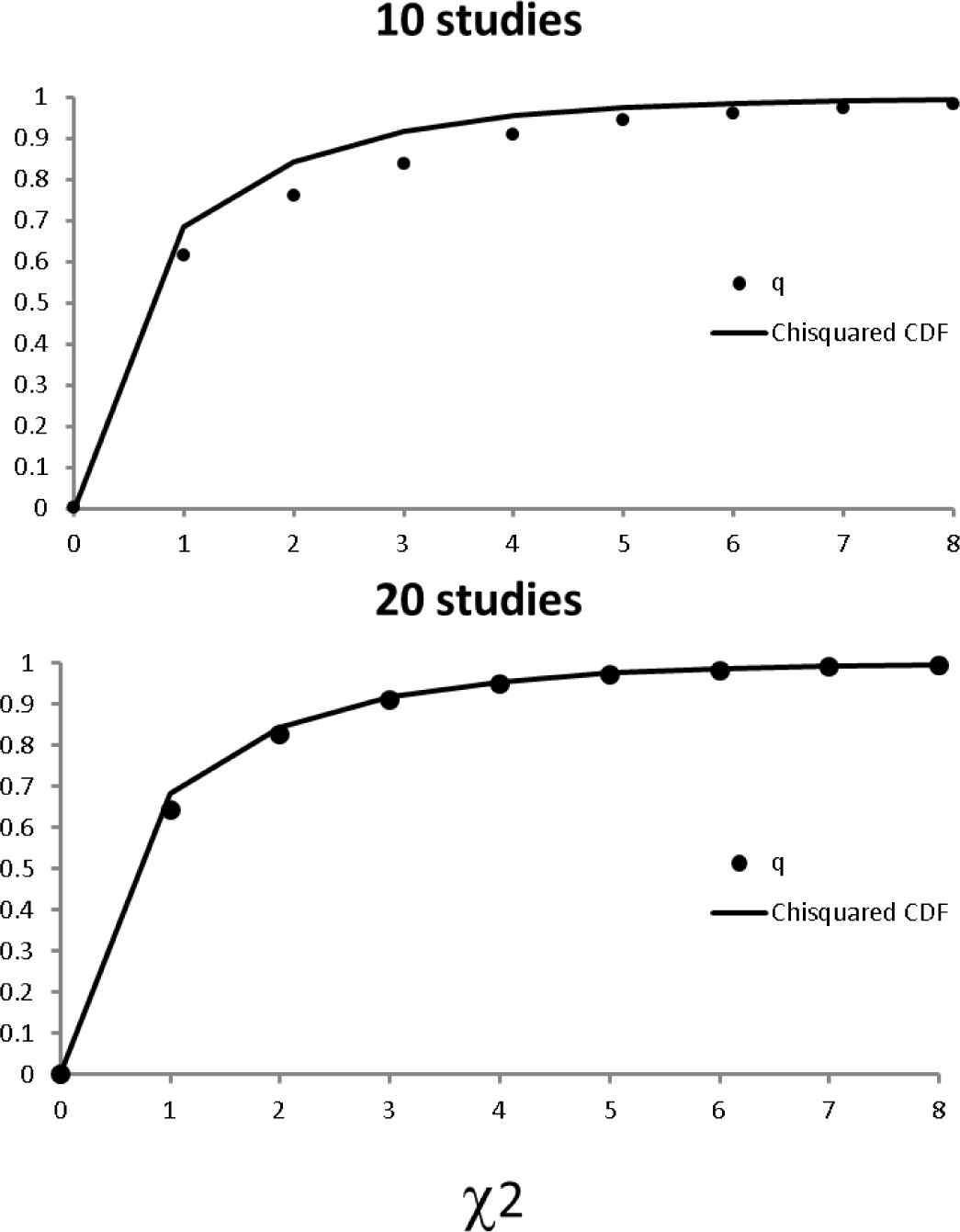
Distribution of test statistic for null (correlation=0) simulated data. Theoretically this is asymptotically (χ^2^ (1 degree of freedom) distributed. The cumulative distribution for the (χ^2^ is shown as a solid line, while simulated results are circle markers. Examples with 10 and 20 simulated studies are presented. Asymptotic convergence to the theoretical distribution is reasonable by 20 studies.

By simulating known networks CBMAN can be shown to detect MVN censored data and threshold the results by FDR. Networks with 6 clusters (nodes) were generated with various MVN covariance matrices. The 6 clusters were reliably detected by CBRES MA for each network variant, and the resulting clusters shown in figure 3.

**Figure 3.**
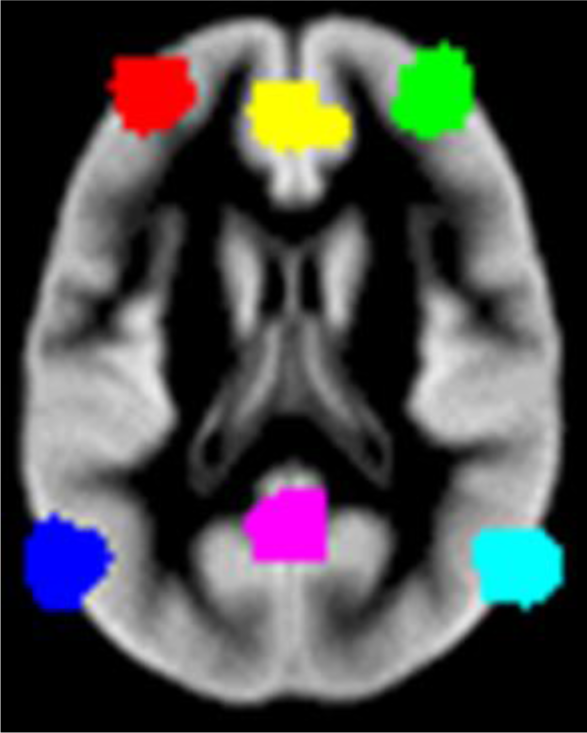
The clusters detected by CBRES MA of the simulated network test data; these results are independent of the network structure.

Figure 4 shows the results of CBMAN on 4 different network configurations that use the 6 clusters shown in figure 3. The network formed by the 6 clusters is determined by the covariance matrix. The first (top left) shows the network with a diagonal covariance matrix where all correlations are statistically zero, hence no network is detected. The second network (top right) includes only one off diagonal element (covariance) and so the network consists only of two nodes connected by one edge. The third example (bottom left) shows a network with a block diagonal covariance matrix, which forms two independent networks as depicted. Finally (bottom right) the covariance matrix specifies a network where effect sizes in each cluster are correlated with every other, as successfully found by CBMAN. Contrasting figure 3 with results shown in figure 4 demonstrates the difference between CBMA and CBMAN; CBMAN considers how the ES correlate between clusters, while CBMA has no mechanism to consider such correlation.

**Figure 4.**
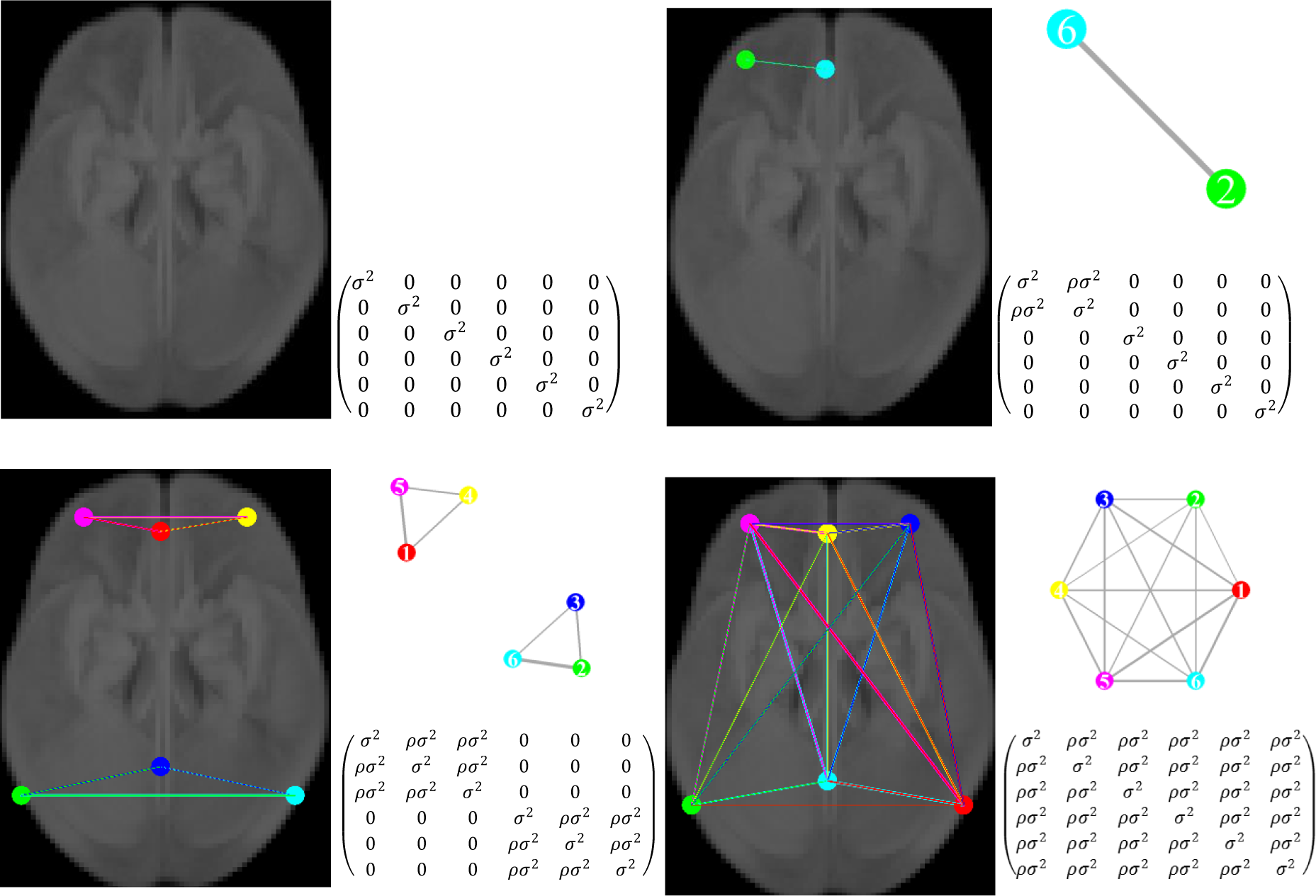
Results of coordinate based meta-analysis of networks. Four networks are simulated using the clusters shown in figure 3. In each case the statistically significant nodes (clusters) and edges are shown, along with the associated graph and the form of the covariance matrix used to generate the multivariate normal effects that define the network.

### Painful thermal stimulus in healthy subjects

Functional MRI studies using painful thermal stimulation of healthy subjects were collected and used to demonstrate CBRES MA. The same data are now used to demonstrate CBMAN. Figure 5 shows the clusters discovered with CBRES MA using a FCDR of 0.05, it also shows the clusters discovered by CBMAN using FDR of 0.05. The significant clusters discovered by both approaches are very similar. The CBMAN analysis shows significant correlations between all clusters. Thresholding the correlation such that ρ^2^>0.5 reduces the number of edges to show only the strongest correlations.

**Figure 5.**
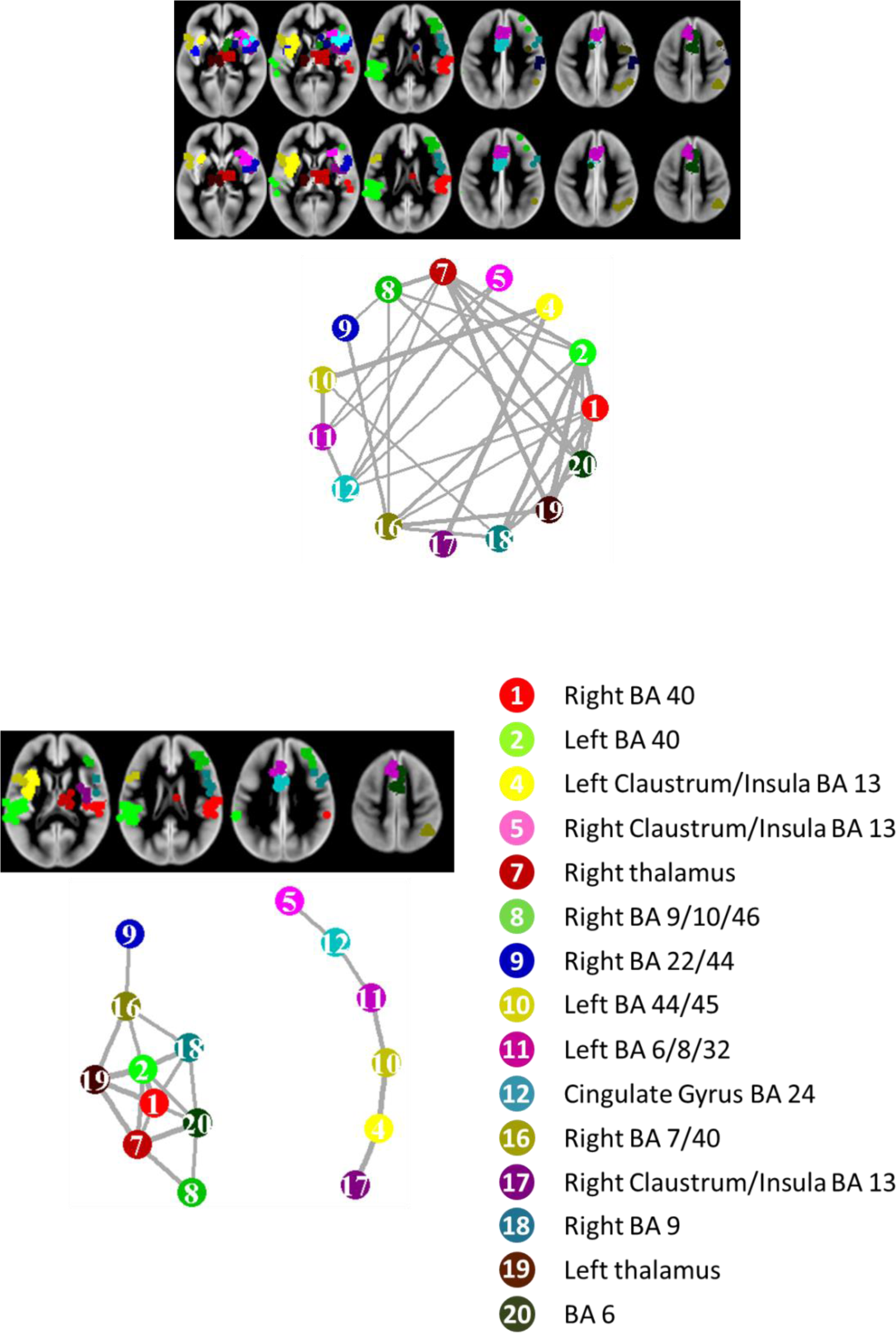
Top: the CBRES MA and CBMAN analysis on functional MRI studies of thermal painful stimulus. The significant clusters produced by CBRES MA are largely similar to those produced by CBMAN at FCDR of 0.05. The graph shows that ES in clusters tend to correlate with each other. Bottom: a correlation threshold of |ρ|>0.707 (ρ^2^>0.5) has been applied reduce the number of edges in the graph, leaving just moderate-strong correlations.

### Voxel based morphometry in Multiple Sclerosis

VBM studies of MS patients were collected to demonstrate CBRES MA, and are used unaltered here to demonstrate CBMAN. Figure 6 shows the clusters declared significant by both methods, and they are similar despite the very different hypotheses tested. For CBRES MA a FCDR of 0.05 was used. For CBMAN a FDR of 0.055 was used because several edges became significant compared to FDR of 0.05. The correlation between symmetric clusters is generally significant.

**Figure 6.**
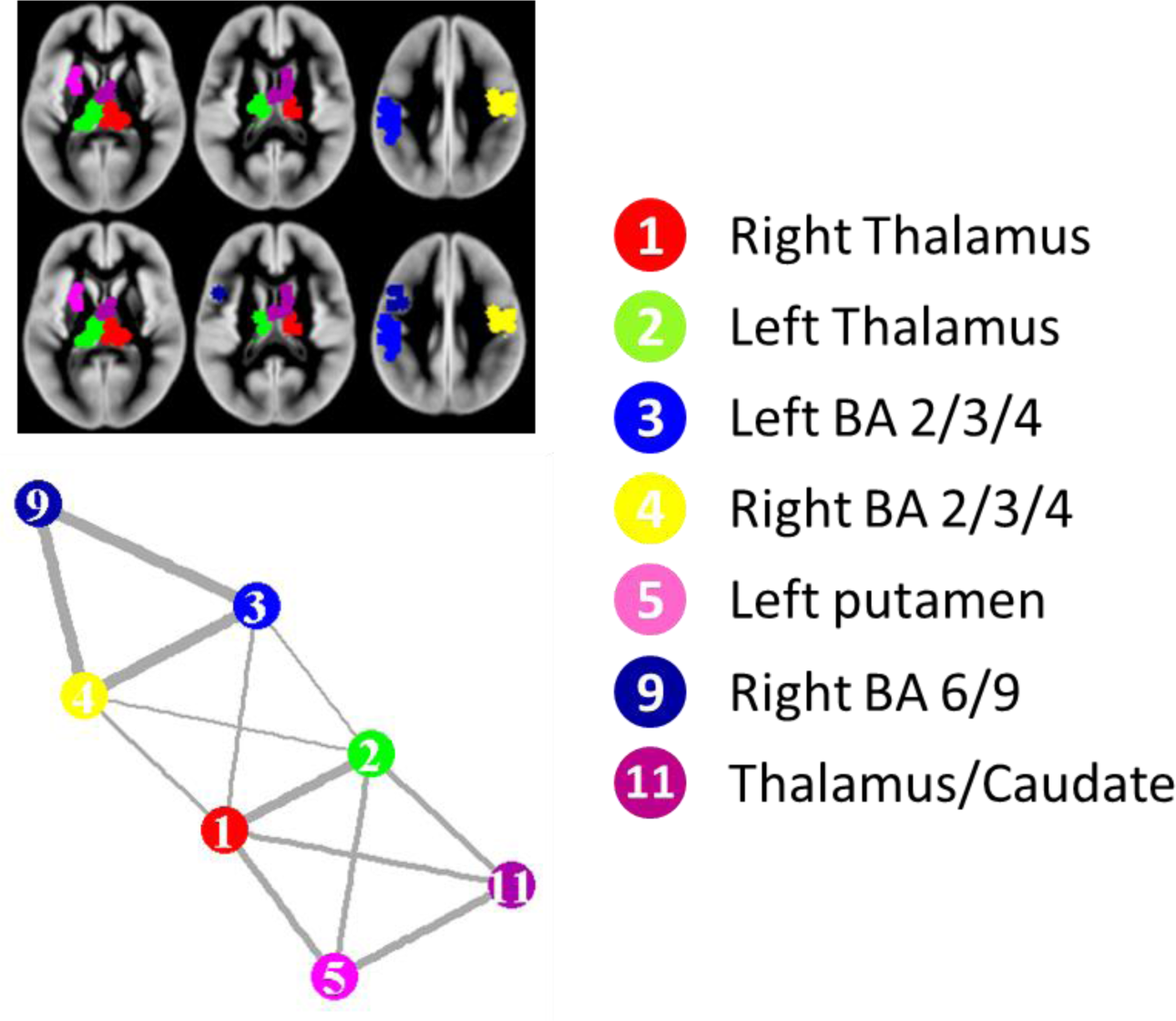
Clusters found by CBRES MA and CBMAN are similar for VBM studies of MS. For the CBMAN analysis an FDR of 0.055 was used as this revealed extra edges. The graph shows strong apparent symmetry between clusters in the two hemispheres.

## Discussion

Here a method of performing a meta-analysis of networks reported by functional MRI or voxel based morphometry studies has been presented. The fundamental assertion is that the multiple effects reported by each study are not independent, but instead represent a single network effect. CBMAN uses only the data needed to perform coordinate based meta-analysis, but tests a very different hypothesis and reveals more features. The results not only show where localised atrophy or activation occurs consistently across studies, but also how the effect is related between the locations.

It is an implicit assumption of CBMA that the results tabulated in fMRI and VBM articles are independent; or at least any dependency is not considered. Yet it is known that spatially separate brain regions act together as a network, rather than in isolation. Multiple activations/grey matter changes might therefore be more appropriately considered as a single related pattern of effect. The pattern is formed by multiple spatially separated clusters with effect-sizes that correlate; in CBMAN this is modelled as a multivariate normal distribution of effect sizes. Fitting the high dimensional MVN distribution to typical neuroimaging summary data is a non-linear problem, however the mathematical property that the marginal bivariate normal distributions involving pairwise clusters have the same parameters as the respective MVN parameters makes fitting feasible in the presence of censored data. Handling censored data when performing maximum likelihood estimation is by integration over portions of the BVN distribution, which is accurately handled numerically.

Coordinate based meta-analysis of networks and coordinate based meta-analysis should be considered complementary. If CBMA reveals multiple significant clusters while CBMAN reveals no clusters, this would be indicative of independent reported ES despite the consistent involvement of regions highlighted by the clusters; a simulated example of this was demonstrated using a network generated by MVN ES with a diagonal covariance matrix (see figure 4). The interpretation of such findings might be that while the CBMA reveals regions frequently implicated by the shared hypothesis of the studies, the effect sizes may vary considerably across subjects and between regions. On the other hand if effect sizes are detectibly correlated between spatially separate regions, CBMAN can reveal a network whether such effects are detected by CBMA or not. In this case correlation of ES across studies suggests that an effect in one region implies an effect in other region(s). Understanding which regions act together, and which act with apparent independence, is a significant extra outcome of the meta-analysis performed by CBMAN.

The fitting of the MVN distribution is by maximum likelihood estimation. Figure 1 indicates the ability to estimate a realistic range of parameters for 20 and 50 studies with simulated censoring; as expected estimation is better with more studies. The thresholding of the estimated correlations is then by statistical hypothesis testing. Pearson’s correlation would be appropriate if the data were not censored, but for censored neuroimaging studies the likelihood ratio test is used and the test statistic is given in equation (8). This is an asymptotic test, but numerical simulation under the null showed that even with as few as 20 studies, the asymptotic distribution assumption is met; see figure 2. Knowing the test statistic under the null is important, since it validates the use of FDR to impose principled control of the type 1 error rate. Beyond this, a default minimum correlation magnitude is applied, although this is user adjustable. The default correlation threshold is that which, if Pearson’s correlation were used, would be detectible with 80% power at a significance level of p<0.05. CBMAN saves correlation matrices, so it might also be possible to analyse weighted graphs [18] rather than binary masks produced by statistical thresholding.

In this report CBMAN has been validated using simulated data with known properties. Beyond validation coordinates from published studies have also been used to demonstrate example output from analysis of real data. The two examples are analyses of functional MRI studies (thermal painful stimulation) and of VBM of grey matter (in multiple sclerosis). This data were originally collected to demonstrate CBRES meta-analysis and have not been altered for CBMAN, emphasizing that no additional data are required than for CBMA. It should be noted that some clusters are different to those reported in [13]. That is because the updated clustering algorithm has partitioned some of the larger clusters. In the two examples the clusters found to be significant are similar for both CBRES MA and CBMAN, despite the different hypotheses. The strongest correlations tended to relate to spatial symmetry in the results, and this was true for both functional and VBM analyses. Further work is underway to add functionality to test differences between networks derived from different groups; for example healthy controls and patients. This involves modifying the parameters of the model to allow the two groups to differ in between-cluster ES correlation. The likelihood ratio test can then be used to test the significance of such differences.

The requirements of performing and reporting CBMAN analysis are similar to those of CBMA. Firstly the method assumes that studies are independent. It is vital that multiple experiments on the same subjects are not considered independent [4, 11] as this will produce a known form of bias common to meta-analysis [19], and consequently reduce the quality of evidence. It is also important to provide the data analysed along with any publication; typically multiple experiments are reported per study and it can be difficult to know which experiments have been included, and therefore to reproduce the analysis, without the data. Provision of data in any meta-analysis is a PRISMA (Preferred Reporting Items for Systematic Reviews and Meta-Analyses) [20] requirement, and only involves inclusion of a small text file. The use of principled control of the type 1 error [21] is also necessary in meta-analysis to provide evidence of effect; uncontrolled type 1 error does not provide any measure of evidence, and would allow apparent significant results even under the null hypothesis. In CBMAN, principled control of the type 1 error is by FDR [14], which should be set appropriately small (FDR≤0.1) to prevent possible excessive null results. For reports the location of the clusters (nodes) are relevant, just as for CBMA. Beyond this the correlation between nodes is important, which is provided as output by CBMAN. Code for producing images of networks, such as those given in figures 4-6, is automatically provided; this calls functions from the iGraph library [22] in R [23].

## Summary and conclusions

Coordinate based meta-analysis is a very popular approach for estimating effects that are spatially consistent over multiple independent neuroimaging studies. To achieve this robust statistical analysis is necessary to for the meta-analysis aim of providing high level evidence of effect. CBMA algorithms use a null hypothesis generated by random coordinates, simulating studies with uncorrelated spatial effects, then extract evidence of true effect by controlling the type 1 error rate in a principled manner. The results reflect the spatial distribution of foci. However, this approach implicitly assumes independence of the multiple effects reported per study. In reality, such results are likely correlated and represent a single network effect. Here, by using the same data needed for CBMA but testing a different hypothesis, it has been shown that this network information can be detected statistically. Provision of not just independent location effects, but also the correlation between the locations without further data is a considerable advantage for CBMAN over other coordinate based meta-analysis methods.

## Appendix Pseudo code for clustering algorithm

### Definitions

***Nstudies***: the number of independent studies.

***Study*_*i*_**: the study to which coordinate *i* belongs.

***MinStudies***: the minimum number of studies overlapping a coordinate in another study needed to form clusters. DBSCAN uses at least the number of dimensions+1. Here 4 is used.

***Overlap*_*i*_**: the number of independent studies reporting coordinates within a distance *MaxCD* of coordinate *i*, including the study to which *i* belongs.

***MaxCD***: the distance such that only 5% of random coordinates have *Overlap*_*i*_≥*MinStudies*

***MinCD***: the minimum allowed clustering distance. This is the smallest clustering distance that prevents a standard cluster (formed from *Nstudies* coordinates and having spatial standard deviation in x, y, and z directions of 3.5mm) fragmenting into more than 1 cluster in 99% of numerical simulations.

***Cluster*_*i*_**: the cluster to which coordinate *i* is assigned; initialised to 0.

***CD*_*i*_**: Clustering distance for coordinate *i*. This is the distance that envelopes *MinStudies* points centred on coordinate *i*. This has a minimum allowed value of *MinCD*.

***D*_*ij*_**: the Euclidean distance between coordinates *i* & *j*.

***Distance*_*i*_**: Shortest path distance between coordinate *i* and cluster peak; initialised to large number.

***Neighbourhoodi*(*d*)**: The coordinates *j* that fall within distance *d* of coordinate *i* and:

- *Overlap*_*j*_≤ *Overlap*_*i*_
- *Distance*_*j*_>*Distance*_*i*_+*D*_*ij*_

### Clustering algorithm

1. *C=0*. Priority queue Q=empty. *Distance*_*i*_=*HUGE_DISTANCE* for all *i*
2. Choose the peak coordinate: the coordinate *i* with maximum *Overlapi and* with *Cluster*_*i*_=*0*
3. If *Overlap*_*i*_*≥MinStudies* Increment *C* *Cluster*_*i*_ *= C* *Distance*_*i*_*=0* Find the *Neighbourhood*(*CD*_*i*_) of *i* and enter these points into Q
4. While Q is not empty: Remove coordinate *j* from Q *Distance*_*j*_=*Distance*_*i*_ *+ D*_*ij*_ If *Overlap*_*j*_≥ 1 *then Cluster*_*j*_=*C* If *Overlap*_*j*_≥ *MinStudies* Find the *Neighbourhood*(*CD*_*j*_) of *j* and enter these points into priority queue Q
5. Repeat from 2 until all coordinates considered for clustering

